# The cold-drought tolerance trade-off in temperate woody plants constrains range size, but not range filling

**DOI:** 10.1101/2021.07.24.453622

**Authors:** Giacomo Puglielli, Enrico Tordoni, Aelys M. Humphreys, Jesse M. Kalwij, Michael J. Hutchings, Lauri Laanisto

## Abstract

Interspecific differences in plant species’ ranges are shaped by complex mechanistic interactions, which have so far remained largely beyond the reach of comprehensive models and explanations. Previous attempts to find underlying mechanisms by examining physiological tolerances to cold and heat separately have yielded contradictory results. Here we test the hypothesis that, instead of examining single stressors, abiotic stress tolerance syndromes that involve trade-offs between multiple abiotic stressors (namely drought, cold, waterlogging and shade), will provide reliable explanations.

We compiled a dataset of actual range size and range filling (the ratio between actual and potential species range) as range metrics for 331 temperate woody plants species from Europe and North America. Tolerance syndromes were expressed as two PCA axes. One axis reflects a drought-cold/waterlogging tolerance trade-off (cold/wet-drought trade-off), the second axis represents a shade tolerance spectrum. Phylogenetic generalized linear mixed models were used to model the range metric-tolerance axes relationships using latitude as an additional main effect, and phylogeny and plant functional type as random effects.

Actual range scaled negatively with the cold/wet-drought tolerance trade-off axis, mostly independently of latitude and continent. Thus, cold/wet-tolerant species had the largest ranges and drought tolerant species the smallest. The sign (−) of the relationship was independent of phylogeny and plant functional type. In contrast, range filling depended on latitude. However, deciduous and evergreen species displayed different distributions of range metrics and tolerance syndromes. No significant relationships with the shade tolerance spectrum were found.

Our findings demonstrate that the cold/wet-drought trade-off partly explains interspecific range size differences. However, this trade-off did not explain range filling. We also showed that fundamental adaptations of species also significantly influence range sizes – stress avoidance through the deciduous habit also explained interspecific differences in range size.

## Introduction

The identification of the factors shaping species range size and filling is a major focus of biogeography (Brown, 1984; Gaston *et al.*, 2009). In recent years, a number of theories aimed at explaining species distribution patterns at the global scale have been developed. For example, the mid-domain effect predicts that species close to the equator have larger distribution ranges (Colwell & Hurtt, 1994), whereas Rapoport’s rule assumes the opposite (Stevens, 1989). Several mechanistic hypotheses either addressing extrinsic factors (e.g. the climate variability hypothesis (Stevens, 1989) to explain Rapoport’s rule) or intrinsic factors (e.g. the dispersal (Hanski *et al.* 1993) or niche breadth (Brown, 1984) hypotheses) have been proposed (reviewed in Sheth *et al.*, 2020). Most of the hypotheses formulated so far to explain interspecific differences in range size have emerged from zoology (Fine, 2015). In comparison, this topic has been little investigated in plant species (Sheth *et al.*, 2020), for which the determinants of range size, and its global variation, remain elusive.

Different metrics of range size are used in the literature to describe plant biogeographic patterns (Sheth *et al.*, 2020). These metrics include latitudinal or longitudinal range, extent of occurrence or area of occupancy, along with the ratio of realized range size (based on known species distributions) over potential range size (usually estimated using species distribution models), also known as range filling (Svenning & Skov, 2004; Paul *et al.*, 2009; Sheth *et al.*, 2020). Despite the use of different metrics, some consistent biogeographic patterns have emerged. For example, North American regions with strong climate instability have lower species richness and large-ranged species, while small-ranged species inhabit more species-rich regions with more stable climates (Morueta-Holme *et al.*, 2013; McFadden *et al.*, 2019). For European tree species, both potential range size and range filling increase with latitude (Svenning & Skov, 2004; Nogués-Bravo *et al.*, 2014), and both European and North American woody species tend to have larger ranges at higher latitudes (Morin & Chuine, 2006).

All these results are consistent with Rapoport’s rule, which predicts larger ranges and lower species richness at higher latitudes, possibly due to high latitude species being more tolerant of more variable environmental conditions (e.g., Morueta-Holme *et al.*, 2013), or to an increased frequency of pioneer species with broader niches at higher latitudes (Morin & Chuine, 2006). Despite these general tendencies, exceptions to Rapoport’s rule have been documented. For example, in the Americas, woody species’ range size has a bimodal distribution in relation to latitude, being largest in both north temperate and tropical areas (Weiser *et al.*, 2007). A similar bimodal distribution has been observed for range filling in European (Svenning & Skov, 2004) and North American tree species spanning sub-tropical to boreal climates (Seliger *et al.*, 2021). Thus, although a positive range size-latitude relationship is well documented for plants, it is not without exceptions and a mechanistic explanation of this pattern is still missing.

Morin & Chuine (2006) proposed that the proximate driver behind Rapoport’s rule is abiotic stress tolerance. In particular, they argued that intrinsic differences in abiotic stress tolerance between species, and thus in their ability to persist under given resource regimes, might explain interspecific variation in range size. Following this proposal, analysis of species’ inherent abilities to withstand extreme heat and/or cold has provided the main way to seek a link between species’ physiology and distribution ranges, chiefly expressed as latitudinal limits (mostly for animals, e.g. Addo-Bediako *et al.*, 2000; Gaston *et al.*, 2009; Sunday *et al.*, 2011; Araújo *et al.*, 2013). Comprehensive large-scale datasets of thermal tolerances have only appeared recently for plant species (e.g., Lancaster & Humphreys, 2020). Nevertheless, relating species’ abiotic stress tolerances to their distribution ranges is complex. Cold and heat tolerance, for instance, only have clear relationships with latitudinal and climatic gradients under certain conditions. Cold tolerance seems to be more closely related to climatic conditions than heat tolerance (Araújo *et al.*, 2013; Lancaster & Humphreys, 2020), and patterns are stronger for northern hemisphere than southern hemisphere species. Similarly, cold and drought tolerances do not display straightforward relationships with range filling, despite showing a clear latitudinal pattern across Europe (Nogués-Bravo *et al.*, 2014). Thus, our comprehension of the relationship between abiotic stress tolerance and species distributions remains poor.

Relating species’ physiological tolerances to their geographical distribution patterns is complicated by occupied ranges representing realized niches (e.g. Hutchinson, 1957), whereas physiological tolerances should reflect species’ fundamental niches. Furthermore, physiological tolerances of different stressors might trade-off against each other due to correlation with independent niche axes of the Hutchinsonian hypervolume (Sexton *et al.*, 2017), shrinking the number of feasible tolerance combinations. Accounting for trade-offs between multiple tolerances may more closely reflect species’ realized physiological requirements (Sack, 2004; Niinemets & Valladares, 2006; Laanisto & Niinemets, 2015; Puglielli *et al.*, 2021a) and possibly reveal consistent relationships with realized range sizes. Thus, multivariate trade-off axes between different tolerances might be needed to detect correlates with range size. Recently, Puglielli *et al.* (2021a) examined multivariate trade-offs in woody species’ ecophysiological tolerances of four major abiotic stresses (cold, shade, drought and waterlogging). The trade-offs were visualized in a triangular stress tolerance space (henceforth *Stress Space*). Ecophysiological tolerance is defined as a species’ ability to survive long-term extreme shortage of a given resource in its natural environment (Niinemets & Valladares, 2006). The *Stress Space* was built using published species-specific tolerance data (Niinemets & Valladares, 2006; Laanisto & Niinemets, 2015) for ~800 Northern Hemisphere woody plant species. Each pair of coordinates in the *Stress Space* reflects species-specific multi-stress tolerance syndromes shaped by trade-offs among the different tolerances. In particular, the first *Stress Space* axis reflects a trade-off between drought and cold/waterlogging tolerance. The second axis is a shade tolerance spectrum, from low- to high shade tolerance. Thus, the *Stress Space* framework permits to link abiotic stress tolerance syndromes to other aspects of species’ biology, including range size. A well-developed framework describing the relationship, and potential trade-offs, between multiple abiotic stress tolerances is crucial for making realistic inferences about the role of abiotic stress tolerance in shaping species distribution patterns, including global variation in range sizes (Gaston *et al.*, 2009).

Identification and interpretation of range size correlates has the potential to increase understanding of the factors and processes that influence species’ ranges (Svenning & Skov, 2004; Estrada *et al.*, 2016). Although a complete spectrum of range size correlates is not yet available (Estrada *et al.*, 2016, 2018), especially for plants, we employ a dataset on 300 temperate woody species from Europe and North America, to analyze the relationships between abiotic stress tolerance syndromes, as summarized by the *Stress Space* axes, and species actual range sizes and range filling. We decided to consider both of these range metrics because they reflect different aspects of species’ ranges. Actual range size includes historical legacies (e.g. for temperate species, the degree to which they have been able to expand since the Last Glacial Maximum; Svenning *et al.*, 2008; Normand *et al.*, 2011; Nogués-Bravo *et al.*, 2014; Estrada *et al.*, 2016) and ecological constraints (e.g. areas with suitable habitat, Linder *et al.*, 2013). Range filling, on the other hand, is a measure of the extent to which species’ ranges are at a climatic equilibrium (Svenning & Skov, 2004). Thus, the traits that are positively correlated with actual range size are expected to be those that are associated with range expansion (Estrada *et al.*, 2016, 2018), whereas traits associated with range filling reflect limits on species’ distributions imposed by historical non-climatic factors (Estrada *et al.*, 2016). Previous studies have mostly attempted to link range filling to plant dispersal syndromes (Svenning *et al.*, 2008; Normand *et al.*, 2011; Nogués-Bravo *et al.*, 2014). However, even assuming successful migration and dispersal, the *regeneration niche* theory (Grubb, 1977), also requires that a species is able to survive the prevailing abiotic (and biotic) conditions to establish viable populations (Estrada *et al.*, 2018). The *Stress Space* framework can therefore provide further insights into the determinants of species’ range size and range filling.

In this study, we examined the relationships between temperate woody species range sizes and range filling, and the *Stress Space* axes. Assuming these metrics are positively correlated (e.g. Seliger *et al.*, 2021), we propose that species’ abiotic stress tolerance syndromes (reflected in their positioning along the *Stress Space* axes) can largely explain latitudinal differences in range size and filling. Specifically, we hypothesized that:

1. As the first *Stress Space* axis represents a spectrum from cold/waterlogging- to drought tolerant species, and assuming this reflects a latitudinal gradient (Nogués-Bravo *et al.*, 2014), we expected a negative relationship between both range metrics and this axis.
2. Both range metrics will be independent of shade tolerance, which in turn is independent of latitude. Shade tolerance data used in the *Stress Space* are measures of a species’ capacity for growth in low light conditions compared to the capacity for growth of coexisting species (Niinemets & Valladares, 2006). Shade tolerance is also largely independent of the other tolerances in the *Stress Space* (Puglielli *et al.*, 2021a). Therefore, shade tolerance can be either high or low irrespective of latitude.

These hypotheses were tested by taking into account the effects of both phylogeny and plant functional type, as defined by Puglielli *et al.* (2021b).

## Methods

### Actual range and occurrence data

We carried out an extensive literature search for polygons defining species’ actual ranges for the 799 woody species in Puglielli *et al.* (2021a). We were able to retrieve polygons defining species’ actual range for 331 species. Specifically, spatial distributions of North American species (*n* = 201) were obtained from the “*Digital representations of tree species range maps from Atlas of United States Trees*” (the digitalized expert-drawn range maps by E.L. Little Volumes 1–5, available at https://github.com/wpetry/USTreeAtlas). According to Seliger *et al.* (2021), Little’s maps are likely the best available approximation of historical realized ranges of North America’s trees and shrubs. This is because E.L. Little intentionally excluded range alterations following Euro-American settlement Seliger *et al.*, 2021). Besides, Seliger *et al.* (2021) also point to the fact that more recent plant occurrence databases are often limited to the continental United States, while many of the tree species we analysed, as for Seliger *et al.* (2021), extend into Canada and Northern Mexico.

Distributions of European species (*n* = 130) were gathered from the International Union for Conservation of Nature (IUCN, www.iucnredlist.org), the European forest genetic resources program (EUFORGEN, http://www.euforgen.org/species), and published papers (Kalwij *et al.*, 2014; Caudullo *et al.*, 2017; Wazen *et al.*, 2020). We consider using expert-based polygons as a proxy of realized range size for European species set a reasonable assumption that is strongly needed to make the results between continental species sets comparable.

Species occurrence records for the 331 species were obtained from the Global Biodiversity Information Facility (GBIF, www.gbif.org/, accessed 21/12/2018; full list of data sources in **Appendix S1**, **Supporting Information Table S1.1**). GBIF data were carefully cleaned using both standardized and customized procedures (see **Appendix S1, Fig. S1.1**).

### Potential range size and range filling calculation

To estimate species’ potential range sizes, we used two presence-only models - i.e., Bioclim (Busby, 1986), and Maxent (Phillips *et al.*, 2006) - to account for differences in potential range size that may arise from algorithmic differences (Nogués-Bravo *et al.*, 2014). Nogués-Bravo et al. (2014) also argued that, giving a possible mismatch between species’ realized and potential (i.e. modeled) species distributions, carefully implemented presence-only methods are expected to return more reliable and parsimonious potential range estimates compared with more complex model classes. Each model was fitted with three environmental parameters: growing degree days at 5°C (GDD, unitless); climatic moisture index (the ratio of annual precipitation to annual potential evapotranspiration, CMI, unitless) and mean minimum temperature of the coldest month (T_min_, °C). GDD and CMI data were obtained from the ENVIREM dataset (Title & Bemmels, 2018) and T_min_ (i.e. Bio6) from WorldClim (Hijmans *et al.*, 2005); each at 10 arcmin resolution. We decided to use only these three environmental predictors for two main reasons: (i) these variables are historically considered as key environmental factors summarizing the processes that limit the spatial distribution of plant species, and they were consistently used in previous studies (e.g. Svenning & Skow, 2004; Nogués-Bravo *et al.*, 2014; Seliger *et al.*, 2021 and references herein); (ii) the use of few environmental predictors carefully selected for their direct effects on plant species distributions avoid artificial and uninformative selection of predictors on the sole basis of statistical techniques (e.g. variance inflation factor).

Before computing Species Distribution Models (SDMs), gathered GBIF occurrence data were subjected to environmental filtering following Varela *et al.* (2014; see **Appendix S1, Fig. S1.1**). All the 331 species had > 20 occurrences, which is considered a reasonable threshold for fitting SDMs (Guisan *et al.*, 2017). The SDMs were fitted using the *sdm* R package (Naimi & Araújo, 2016). For each run, 80% of species data was used for training, and the remaining 20% for evaluating the model. 30 replicates per species were generated through bootstrapping and 20,000 background points were generated at each run. The Area under the ROC Curve (AUC; Fielding & Bell, 1997) and True Skill Statistic (TSS; Allouche *et al.*, 2006) were used to evaluate model performance. The SDMs predictions were converted into presence/absence maps by using the threshold that maximized both sensitivity and specificity of the model. This is considered the best option for presence-only methods (Liu *et al.*, 2013). The number of suitable 10 arcmin cells in the binary maps corresponded to potential range size while actual range was determined by counting the 10 arcmin cells occupied by the polygons defining species’ actual range. Range filling (%) was then calculated as: (Actual range/Potential range)×100 (see **Appendix S2, Fig. S2.2**). Due to a lower percentage of species with range filling estimates greater than 100% (see **Appendix S3, Figs. S3.3-3.6, Table S3.2** for considerations on models’ performance and range filling estimates) only Bioclim derived estimates of potential range were used in subsequent analyses, as in Nogués-Bravo *et al.* (2014). Furthermore, as pointed out by the same authors, similar studies of range filling (e.g. Svenning & Skov, 2004) also used the Bioclim algorithm to estimate potential range size for range filling calculation, thus ensuring that our results are broadly comparable across a broader literature set.

In order to account for broad differences in species’ adaptive syndromes, species were classified according to three major plant functional types: deciduous angiosperms, evergreen angiosperms and evergreen gymnosperms. For the complete list of species’ actual range size (log_10_-transformed number of 10 arcmin cells), range filling, centroid latitude and species classification according to their continental origin (N. America, Europe) and plant functional type see **Appendix S4**.

### Abiotic stress tolerance data

The species-specific estimates of tolerance of shade, drought, cold and waterlogging used to define the *Stress Space* were obtained from the datasets of Niinemets & Valladares (2006) and Laanisto & Niinemets (2015), which include stress tolerance scores for ~800 Northern Hemisphere woody species. In the original data compilation (Niinemets & Valladares, 2006), shade, drought and waterlogging tolerance were independently estimated by cross-calibrating multiple tolerance scales reported in the literature where multiple measurements for one species were available across tolerance scales. Cold tolerance data were extracted from USDA plant hardiness data and represent species-specific averages gathered from multiple sources (see Laanisto & Niinemets, 2015 for further details). In this respect, cold tolerance data do not show any circularity with T_min_ obtained from WorldClim as it is not based on any spatially-explicit information, and being an average species-specific estimate, it is not affected by a species range size (i.e. species with larger ranges are more likely to encounter extreme mean temperatures). All the stress tolerance scores vary in a continuous fashion between 1 - very intolerant species - to 5 - very tolerant species (Niinemets & Valladares 2006 and Laanisto & Niinemets 2015).

The formalization of the *Stress Space* (Puglielli *et al.*, 2021a) revealed that two-dimensions (principal components) capture ~80% of the variance in species-specific combinations of shade, drought, cold and waterlogging. Each pair of coordinates in the *Stress Space* corresponds to a species-specific stress tolerance syndrome. Stress Axis 1 is positively correlated with drought tolerance and negatively correlated with both waterlogging and cold tolerance. It is interpreted here as a cold-drought tolerance trade-off, where the term cold refers to a short growing season. This interpretation stems from the positive covariance between cold and waterlogging tolerance in our dataset: the highest cold tolerance is expected where snowpacks are greater, resulting in later snowmelt, followed by waterlogging and consequently a shorter growing season (Chuine, 2010). Stress Axis 2 is positively correlated with shade tolerance, and represents a shade tolerance spectrum. Stress Axes are available in Puglielli *et al.* (2021a).

### Data analysis

Actual range size (hereafter range size) was log-transformed before analysis. Range filling was strongly related with range size (*R^2^* = 0.46, p < 0.0001, *n* = 331), but not with log_10_-transformed potential range size (*R^2^* = 0.02, p < 0.001, *n* = 331). Thus, we used the residuals deriving from the relationship range size vs. range filling as a metric of range filling (Seliger *et al.*, 2021).

The relationships between stress axes and both range size and range filling residuals were tested at different levels. First, we used Ordinary Least Square (OLS) regression analysis. Second, we used quantile regression to provide a more comprehensive characterization of the studied relationship (Ricotta *et al.*, 2010). In addition, quantile regression is mostly insensitive to outliers (Ricotta *et al.*, 2010). Quantile regressions were run using the *quantreg* R package (Koenker, 2017) using different quantiles of range metrics distribution (τ = 0.1, τ = 0.25, τ = 0.50, τ = 0.75 and τ = 0.90). However, fitting models with many species while ignoring phylogenetic relationships might lead to inflated type I errors (Freckleton *et al.*, 2002). Consequently, as a third step, we computed Phylogenetic Generalized Linear Mixed Models (PGLMMs, Ives & Helmus, 2011) using range metrics as response variable as a function of stress axis 1 or 2 (considered separately), centroid latitude obtained from cleaned GBIF occurrence data using the *geosphere* R package (Hijmans *et al.*, 2019), plus the first-order interaction stress axis : latitude; species’ phylogenetic relatedness and plant functional types were considered as random effects. When the interaction was not found to be significant, only the main effects were considered. PGLMMs were run using the *phyr* R package (Li *et al.*, 2020). Phylogenetic data were retrieved for 325 species in our dataset using the mega-tree available via the *V.PhyloMaker* R package (Jin & Qian, 2019). The mega-tree combines the phylogenies developed by Zanne *et al.* (2014) and Smith & Brown (2018). Species nomenclature followed The Plant List v.1.1 (2013). We also tested for potential signals of spatial autocorrelation in the model residuals using spline correlograms from the *ncf* R package (Bjørnstad, 2020); specifically, 95% pointwise bootstrap confidence intervals were computed from 5,000 bootstrap samples of Pearson residuals.

Finally, we tested for differences between plant functional types in terms of the distributions of actual range and range filling residuals, and positioning along stress axes using the Kruskal-Wallis test. Pairwise multiple comparisons between group levels were carried using Dunn’s test by adjusting p-values with Holm correction.

All the data analysis procedures were repeated for the European and North American species separately to account for possible geographic differences in patterns observed. All statistical analyses were performed in R 4.0.5 (R Core Team, 2021). As we did not find any relationship between either range size or range filling residuals with the shade tolerance spectrum at any level of analysis, only the results relative to the cold-drought tolerance trade-off axis are shown.

## Results

Actual range scaled negatively with species positioning along the cold-drought tolerance trade-off axis for species from both continents (Europe: slope = −0.10, *R*^2^ = 0.06, *p* = 0.01, *n* = 130; North America: slope = −0.18, *R*^2^ = 0.14, *p* < 0.01, *n* = 201) (**Fig. 1 a,c**). A negative relationship between actual range and cold-drought tolerance trade-off axis was also observed across the considered quantiles of the response variable, but with differences between the species from the two continents. For European species, the quantile regressions were mostly significant at average to high values of actual range (**Fig. 1 a**). For North American species quantile regressions were all significant, except the one fitted at the lowest quantile (**Fig. 1 c**).

**Fig. 1.**
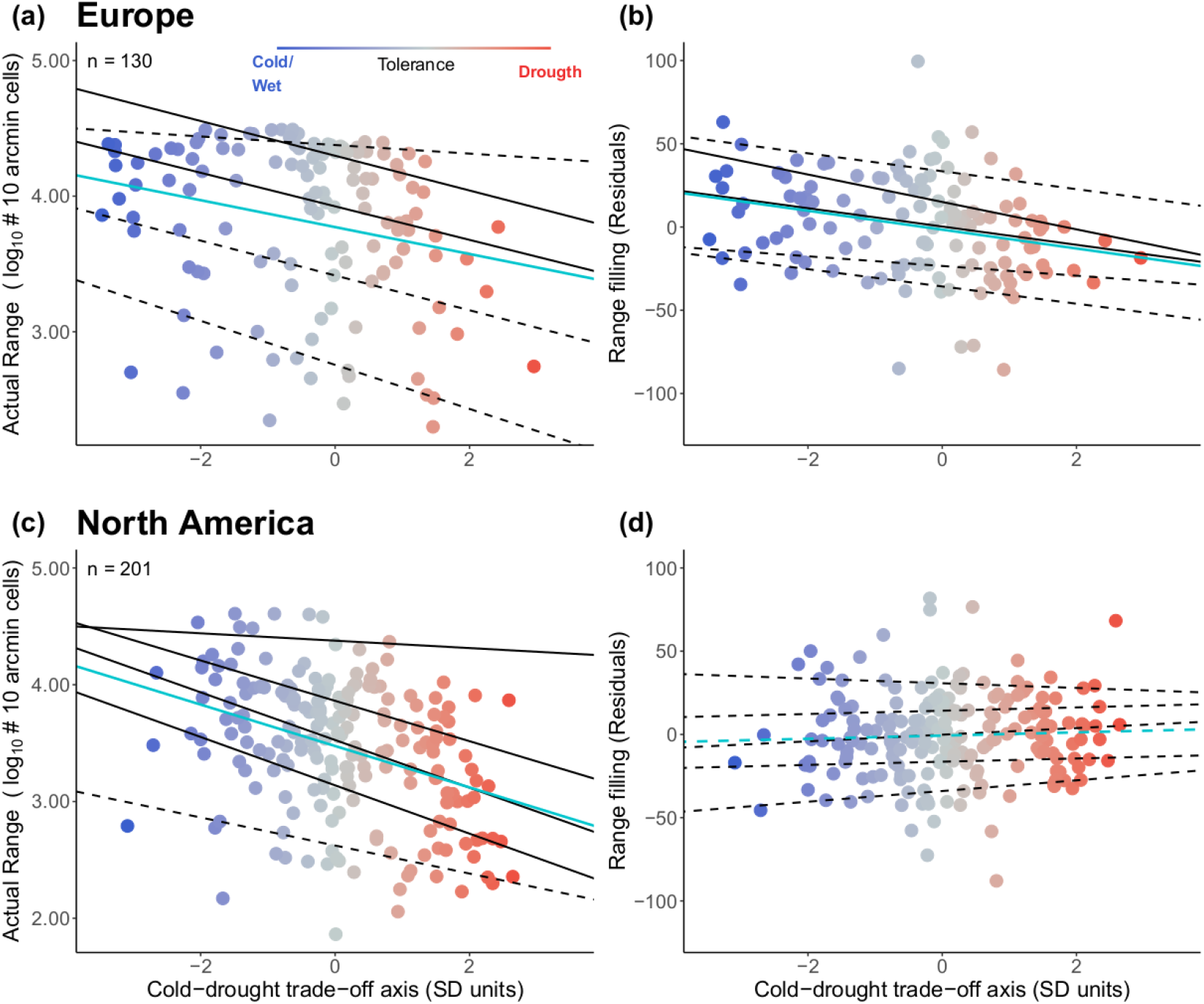
Relationship between the cold-drought trade-off (Standard Deviation units) and actual range (log_10_-transformed number of 10 arcmin cells) and range filling (residuals, see Methods) for: **(a,b)** European, and **(c, d)** North American woody plant species. Pale blue line represents the Ordinary Least Square fit. Black lines show the quantile regressions fitted at the 0.1, 0.25, 0.50, 0.75 and 0.90 quantile of the response variable distribution. Solid lines depict significant relationships at p < 0.05 while dashed lines represented not significant relationships. Sample size (n) is shown in panels **(a,c)** and applies also to the relationships involving range filling as the response variable. The color gradient reflects the progression from cold/wet-tolerant to dry/warm-tolerant species along the cold-drought trade-off.

Range filling residuals scaled negatively with species positioning along the cold-drought tolerance trade-off axis only for European species (slope = −5.66, *R*^2^ = 0.07, *p* < 0.01, *n* = 130) and, as for actual range, the quantile regressions were mostly significant at average to high values of range filling (**Fig. 1 b**). No significant relationship was found between range filling residuals and the cold-drought tolerance trade-off axis for North American species, and the relationship was not significant at any considered quantile (**Fig. 1 d**).

The negative relationship between actual range and the cold-drought tolerance trade-off axis was not affected by including latitude as an additional main effect (**Table 1**), and the cold-drought tolerance trade-off axis effects were always greater than that of latitude for all data pooled (**see Appendix S5, Table S5.3**), and for each continent considered separately (**Table 1**). However, some differences between continents were detected. The cold-drought tolerance trade-off axis effect was only marginally significant (p = 0.06) for European species and the effect of latitude was not significant. The model explained 7% of the variance of actual range for European species. In contrast, for North American species, the effects of both the cold-drought tolerance trade-off axis and latitude were significant, but with opposite sign: the tolerance axis maintained its negative relationship with actual size whereas latitude showed a positive relationship (p < 0.01 and p < 0.05, respectively). This model explained 23% of the total variation in actual range of North American species, and indicates that range sizes are greatest at higher levels of cold/wet tolerance and at higher latitudes. Despite European and North American species sets having no species in common, we explored whether differences between continental species sets could be driven by differences in terms of genera composition.

**Table 1.** Results of the Phylogenetic Generalized Linear Mixed models.

| Response | Continent | Main effects |  |  |  |  |  | Random effects |  |
| --- | --- | --- | --- | --- | --- | --- | --- | --- | --- |
|  |  |  | Estimate | SE | *p* | Model *R*<sup>2</sup> | *n* |  | Variance |
| Actual range | Europe | Intercept | 3.39 | 0.34 | < 0.0001 | 0.07 | 128 | PFT | 0.007 |
|  |  | Cold-drought trade-off | -0.07 | 0.04 | 0.06 |  |  | Phylogeny | ≈ 0 |
|  |  | Latitude | 0.01 | 0.01 | ns |  |  |  |  |
|  | North America | Intercept | 2.9 | 0.22 | < 0.0001 | 0.23 | 197 | PFT | 0.05 |
|  |  | Cold-drought trade-off | -0.11 | 0.03 | < 0.001 |  |  | Phylogeny | ≈ 0 |
|  |  | Latitude | 0.01 | 0.01 | < 0.01 |  |  |  |  |
| Range filling | Europe | Intercept | 27.08 | 17.37 | ns | 0.19 | 128 | PFT | 33.12 |
|  |  | Cold-drought trade-off | -43.18 | 11.77 | < 0.0001 |  |  | Phylogeny | 0.003 |
|  |  | Latitude | -0.56 | 0.34 | ns |  |  |  |  |
|  |  | Cold-drought trade-off * Latitude | 0.69 | 0.22 | < 0.001 |  |  |  |  |
|  | North America | Intercept | 22.96 | 9.25 | 0.05 | 0.04 | 197 | PFT | 3.97 |
|  |  | Cold-drought trade-off | 0.01 | 1.56 | ns |  |  | Phylogeny | 0.001 |
|  |  | Latitude | -0.6 | 0.22 | < 0.001 |  |  |  |  |

Models were run by continent (Europe and North America) using actual range (log10-transformed number of 10 arcmin cells) and range filling (Residuals, see Methods) as response variables, Cold-drought trade-off (Standard Deviation units) and latitude (centroid latitude, °) as main effects, and Plant Functional Type (PFT) and species phylogenetic relatedness (Phylogeny) as random effects. Estimates of the main effects and their standard errors (SE), together with the variance explained by the model (Model *R*<sup>2</sup>) and sample size (*n*), are shown. The variance captured by the random effects is also shown.

Out of 106 genera in our entire dataset, there were 29 genera in common between the two continents. These genera contributed 70% of the species included in the entire dataset. When the analyses were repeated after removing genera that were unique to one or other continent, the result remained significant for North American species, but the relationship between range size and the cold-drought tolerance trade-off axis changed from being marginally significant to become highly significant for European species as well (R^2^ = 0.12, see **Appendix S5, Table S5.4**). This indicates that differences in results between continents are partly driven by differences in genera composition of their constituent species.

There was no significant interaction between the cold-drought tolerance trade-off axis and latitude in any model involving actual range as the response variable for European or North American species. Overall, species positioning along Stress Axis 1 was the main driver of interspecific differences in actual range for both continents. Actual range data in relation to the cold-drought tolerance trade-off axis and latitude are shown in **Appendix S5, Fig. S5.7a,c.**

The differences in results between continents were more pronounced for range filling (**Table 1**). A significant positive interaction between the cold-drought tolerance trade-off axis and Latitude was found in the model including range filling for European species, and it explained approximately 19% of the variance. Despite a significant main effect of the cold-drought tolerance trade-off axis on range filling (**Table 1**), we did not interpret this effect given the presence of a significant interaction. We regard the interaction between the cold-drought tolerance trade-off axis and latitude as the main driver of interspecific differences in range filling across European species. For North American species, range filling showed a negative relationship with latitude and no significant effect of the cold-drought tolerance trade-off axis. This model suggests greater range filling at lower latitudes, but it explained only 4% of variance, leaving range filling largely unexplained for North American species. Range filling data in relation to the cold-drought tolerance trade-off axis and latitude are shown in **Appendix S5, Fig. S5.7b,d.**

Regardless of the model, the effect of phylogenetic relatedness between species was negligible (**Table 1**). In addition, the spline correlograms (see **Appendix S6, Fig. S6.8**) did not reveal any evidence of spatial autocorrelation in the PGLMMs residuals. We can therefore safely disregard spatial autocorrelation as a factor influencing model parameter estimates. Plant functional types did not affect the strength or the sign of the relationships between the cold-drought tolerance trade-off axis and either actual range or range filling. However, as a random effect in PGLMMs, plant functional type had a greater effect overall than species phylogenetic relatedness in terms of random effect variance (**Table 1**).

The three plant functional types differed in distributions of actual range and range filling values and in positioning along the cold-drought tolerance trade-off axis (Kruskal-Wallis test, p ≤0.05; **Fig. 2 a-f**). As a general trend, deciduous angiosperms have larger actual range and range filling values (p ≤ 0.05; **Fig. 2 a,b,d,e**), and they occupy the cold/wet side of the cold-drought tolerance trade-off (i.e. more negative values along the cold-drought tolerance trade-off axis) (**Fig. 2 c,f**), compared to the other plant functional types. However, multiple comparisons sometimes differed between continents (**Fig. 2 a-f**).

**Fig. 2.**
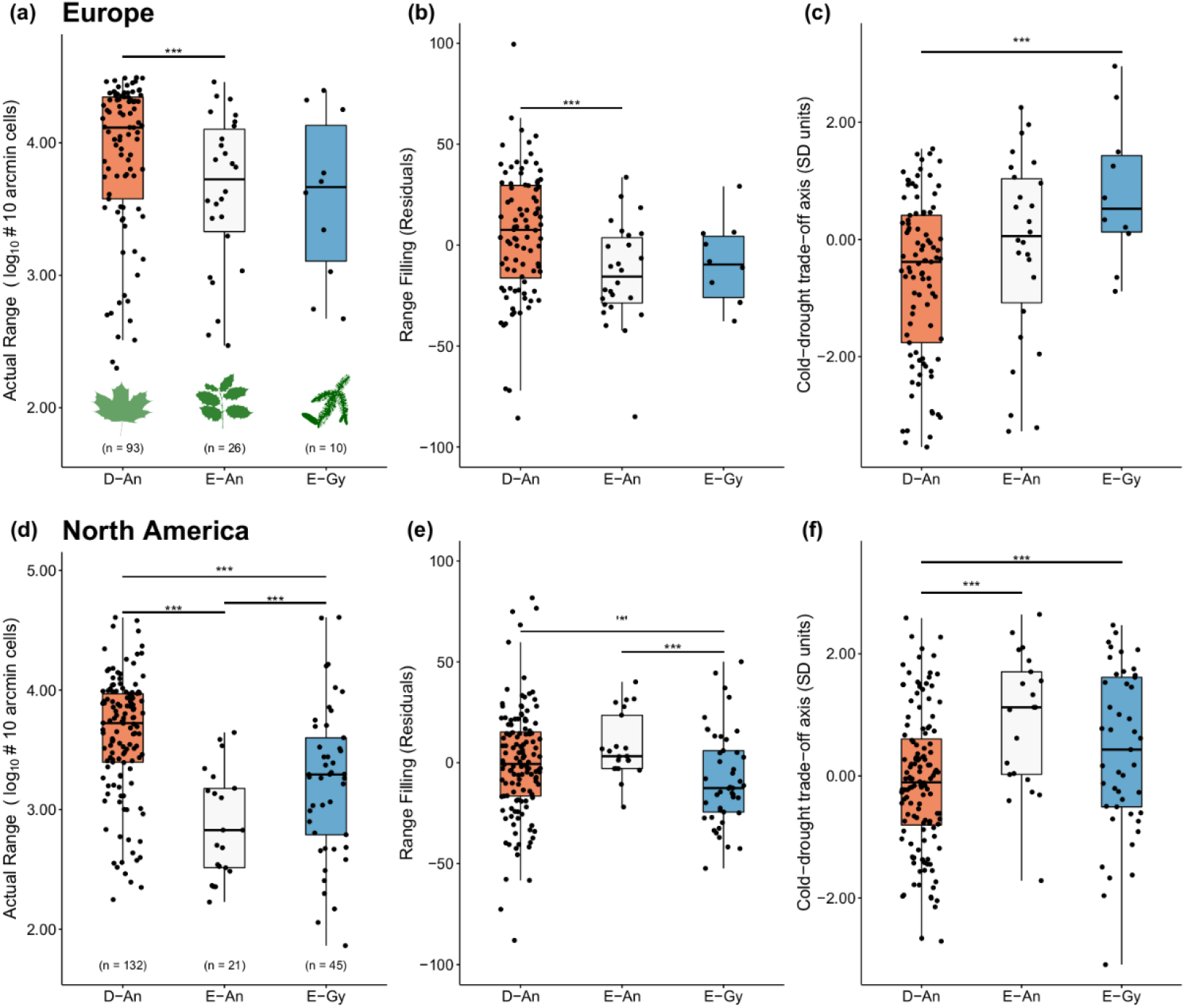
Boxplots for the distribution of actual range size (log_10_-transformed number of 10 arcmin cells), range filling (Residuals, see Methods) and positioning of species along the cold-drought trade-off (Standard Deviation units) within each of the plant functional types (D-An = deciduous angiosperms; E-An = evergreen angiosperms; E-Gy = evergreen gymnosperms) for: **(a-c)** European, and **(d-f)** North American woody species. Sample size and data points for each plant functional type are shown. *** indicates significant differences between plant functional types (Kruskal-Wallis, p < 0.05). ‘*’ indicates marginal significance (p = 0.05). Only significant and marginally significant differences are shown. Multiple comparisons between groups were carried out using the Dunn’s test and Holm correction for multiple testing.

## Discussion

Our results show that the cold-drought tolerance trade-off axis (hereafter cold-drought trade-off) partly explains interspecific differences in actual range size of temperate woody plants (**Fig. 1**). In particular, we found large-ranged species at the cold/wet tolerance end of the trade-off axis, and small-ranged species at the drought tolerance end. Despite some nuanced variation discussed below, this result was independent of continent, latitude, plant functional type and phylogeny, indicating its generality. In contrast, range filling showed different patterns in Europe and North America: the interaction between the cold-drought trade-off and latitude had the largest influence on range filling for European species, whereas latitude was the only driver of range filling in our North American species pool.

### Determinants of range size variation in temperate woody plants

For North American species, we found that cold/wet tolerant species occurring at high latitudes (e.g. *Salix* spp., *Larix laricina* (Du Roi) K.Koch) generally have the largest ranges (**Table 1**). Consistent with this, large ranged, cold-tolerant North American tree species are known to be generally absent from regions that are consistently warm and moist, such as the southeastern regions of the continent (Pither, 2003). Palaeoecological records also provide evidence for rapid northward range shifts in North American large-ranged trees after the latest ice age (Seliger *et al.*, 2021, and references herein). Conversely, some species might have maintained relatively small realized ranges following deglaciation, perhaps due to trait syndromes guaranteeing competitive advantages only in specific ice age refugia (Seliger *et al.*, 2021). Possible examples are species with drought tolerance strategies that prevented northward range expansions after glacial retreat, such as *Juniperus deppeana* Steud., *Pinus monophylla* Torr. & Frém., and *Quercus douglasii* Hook. & Arn., among other species with relatively small ranges that are confined to SW-North America. Long-term drought, and likely adaptations to tolerate such conditions, are an important constraint on plant species distributions (e.g. Normand *et al.*, 2009). Similarly, Pither (2003) hypothesized that latitudinal patterns in the range sizes of North America woody species reflect a potential trade-off between species’ cold tolerance strategies and their competitiveness in warmer environments. In support of this, our results show that the cold-drought trade-off, which largely reflect species biogeographical history, is a mechanism that actively influences interspecific differences in range size for North American plant species.

Cold-tolerant European species also have large ranges (**Fig. 1 a**), but the signal was weaker than for North American species (**Table 1**). This difference is caused by differences in genera composition between the continental species sets (see Results section), and by a cluster of European species with intermediate positions along the cold-drought trade-off having large ranges (**Fig. 1 a**). These large-ranged species include *Picea abies* (L.) H.Karst, *Pinus sylvestris* L. and *Betula pendula*. Roth. Such species survived the last glacial maximum in central and/or eastern Europe with easy access to Northern Europe, with possibility for rapid northward expansion, after ice retreat (Normand *et al.*, 2011). While these species are indeed cold tolerant, their tolerance syndromes include some degree of drought tolerance as well, shifting them towards the center of the cold-drought trade-off axis. This effect has the potential to decouple these species from the negative relationship between the cold-drought trade-off and range size, indicating the importance of considering stress tolerance syndromes based on several factors rather than single tolerances when investigating interspecific differences in ranges. Other species that are potentially decoupled from this relationship belong to European genera that do not occur in the North American species set. More studies are needed for these European species. In the case of small range, drought-tolerant species, the explanation provided for North American species is likely to apply to European species as well (e.g. Normand *et al.*, 2009).

The negative scaling of range size along the cold-drought tolerance trade-off is also consistent with macrophysiological evidence suggesting that pre-adaptation to low temperature or species-specific abilities to adapt to freezing temperatures may have favoured species’ northward migration after glacial retreat (e.g. Araújo *et al.*, 2013; Lancaster & Humphreys, 2020). In contrast, it has been suggested that heat tolerances might be less adaptable (e.g. Araújo *et al.*, 2013) and, considering that drought-tolerant species usually thrive in hot environments, adaptations to tolerate drought might have concomitantly prevented the northward post-glacial race for drought tolerant plants. This suggests that the cold-drought trade-off might limit niche expansion for temperate woody species, as has been suggested for heat stress globally (Araújo *et al.*, 2013). Importantly, however, despite the fact that drought and heat stress can covary in warm-temperate ecosystems, we argue that the effect of drought tolerance on range size should be considered separately from that of heat tolerance. The most heat tolerant plants are from both dry and moist environments (Lancaster & Humphreys 2020), and although transpirational cooling generally costs moisture, plants have also evolved adaptations to reduce the detrimental effects of heat stress under drought (Flexas *et al.*, 2014). For example, among a plethora of other adaptations, changes in water-use efficiency (Flexas *et al.*, 2014) or leaf movements (Puglielli *et al.*, 2017) can decouple leaf physiological responses to high temperatures from those to drought in Mediterranean woody plants.

We have shown that species positioning along the cold-drought trade-off axis imposes fundamental constraints upon interspecific differences in the range size of woody species, and contributes to shaping their realized niches (i.e. actual range) across continents. Species positioning in the *Stress Space*, which is determined by trade-offs between multiple tolerances (Puglielli *et al.*, 2021a), can be interpreted as a measure of a species’ realized niche based on abiotic resources (*sensu* Hutchinson, 1957) more than as a measure of the fundamental niche. After all the trade-off between tolerances shrinks fundamental niche size towards that of the realized niche (e.g. Sack, 2004). Therefore, by representing the typical multiple abiotic stress levels to which a plant species has adapted, its position in the *Stress Space* inherently corresponds to a set of energy constraints that define plant life history strategies. Such strategies are thought to directly control species’ geographical distributions (Morin & Chuine, 2006). Niche position along resource-defined axes is a strong predictor of range size and occupancy in many animal groups (Seliger *et al.*, 2021), and our results show that this is also true for temperate woody plant species.

### Range filling is not affected by the cold-drought tolerance trade-off

All species are expected to have constraints on the extent to which they fill their range because of their specific physiological and ecological requirements (Paul *et al.*, 2009), and their inherent trade-offs. Our results suggest that other factors that covary with latitude, not considered in this study (e.g. dispersal syndromes, Estrada *et al.*, 2016), might be important for explaining range filling pattern. Such historical non-climatic limitations can ultimately also covary with species’ abiotic stress tolerances. This might explain why a positive interaction between the cold-drought tolerance trade-off and latitude drives their range filling differences. Similarly, Nogués-Bravo *et al.* (2014) proposed that the negative correlation between seed mass and range filling found across 38 European tree species could be due to covariation between seed mass and other factors (e.g. drought tolerance), but further analysis including a larger number of species would be needed in order to test this claim.

More importantly, our results highlight different range filling determinants between the two continents, that contrarily to actual range, did not depend on differences in genera between continents (see **Appendix S5**, **Table S5.4**). European species’ range filling was driven by a positive interaction between latitude and the cold-drought tolerance trade-off, while North American species’ range filling was influenced only by a negative effect of latitude. This negative relationship between range filling and latitude for North American species was in agreement with a previous study that also reported a positive relationship range filling-longitude (Seliger *et al.*, 2021), suggesting that longitude can indeed alter the expected range filling-latitude relationship. Differences between continents might therefore depend on their geographical extent. Europe has a much smaller latitudinal and longitudinal range and less gradual geographical clines than North America (Morin & Chuine, 2006). In addition, some European species ranges also occur outside Europe (e.g., species with a Eurasian distributions) and this might have influenced model’s projections (see **Appendix S3**). According to the above, we therefore suggest that the interaction between these contrasting continental features and intrinsic drivers, such as dispersal syndromes, might determine continent-specific levels of range filling.

### Differences among plant functional types

Plant functional type (PFTs) did not affect the negative scaling of range size with the cold-drought trade-off (**Table 1**). However, the distribution of range size, range filling and positioning along the cold-drought trade-off axis differed between PFTs (**Fig. 2 a-f**). This probably explains why PFTs generally accounted for more of the random effects variance (even though the amount was generally low) for range metrics compared to phylogeny (**Table 1**). In general, despite some overlap between PFTs, deciduous angiosperms showed the highest values for both actual range and range filling (consistent with large-ranged species also having greater range filling, Seliger *et al.*, 2021), and they were located further towards the cold tolerance end of the trade-off axis. Using a dataset of European and North American woody species, Morin & Chuine (2006) also found that deciduous species were overrepresented among the large-ranged species.

Many deciduous angiosperms differ in trait syndromes from evergreen angiosperms or evergreen gymnosperms (e.g. Puglielli *et al.*, 2021b), and different trait syndromes match differences in PFTs geographical distributions (Zanne *et al.*, 2018). For example, in the northern hemisphere, deciduous species tend to be more frequent in cold climates compared to evergreen broad-leaved angiosperms, despite some overlap between PFTs at almost all latitudes (Zanne *et al.*, 2018). Conversely, while adaptations to tolerate drought closely match the distribution of evergreen species, this is not always true for deciduous species (e.g. see Kunert *et al.*, 2021), suggesting that drought does not always limit deciduous species spatial distribution.

Deciduousness *per se* has been the common explanation for the ability of deciduous species to colonize either cold or dry environments, as it is an adaptation that permits avoidance of unfavorable environmental conditions. This explains why deciduous angiosperms in our dataset have larger ranges than other PFTs, and are mostly cold-tolerant. However, if deciduousness is also a successful drought avoidance strategy that could drive to large ranges in drought tolerant species as well, the relationship between range size and the trade-off axis should be not significant rather than negative. We can reconcile this by considering that woody species of the warm-temperate/temperate regions of the Northern Hemisphere are not generally drought-deciduous. Drought tolerant deciduous species have sufficiently though leaves to widen their growing season beyond the usual duration of periods of drought (Hallik et al. 2009). Such adaptation is of course part of a wider trait syndrome (see Hallik *et al.*, 2009), including, for example, greater biomass allocation to roots in species with greater drought-tolerance (Puglielli *et al.*, 2021b), or smaller vascular conduits (Olson *et al.*, 2018), that support the investment in tough leaves. This is consistent with the previous discussion of a link between species’ tolerance strategies, trait syndromes and range size.

To summarize, while we recognize that adaptations to very low temperatures are complex, and involve a combination of avoidance and tolerance strategies together with acclimation (Schubert *et al.*, 2020), we argue that large range sizes at the cold tolerant end of the trade-off axis is made viable because of the deciduous habit in temperate woody species. In this instance, stress avoidance is more important as an asset than stress tolerance. However, we also observed a greater overlap, in both range size and positioning along the trade-off axis, between freezing-tolerant North American gymnosperms and deciduous angiosperms than with evergreen angiosperms (**Fig. 2 d,f**). Thus, we do not exclude the additional role for freezing tolerance in guaranteeing large ranges.

## Conclusions

Our results demonstrate that the cold-drought tolerance trade-off partly explains interspecific differences in range size across temperate woody plant species and that this relationship is largely independent of latitude and consistent with woody species biogeographical histories in the considered continents. Notably, our findings also suggest that accounting for species’ abiotic stress tolerance towards multiple stresses can reconcile macroecological and macrophysiological theories aimed at explaining range size differences among woody plant species, supporting a previous hypothesis by Morin & Chuine (2006). However, our results concerning the impact of abiotic stress tolerance on range filling were inconclusive, suggesting that other factors not studied here and that covary with latitude and/or abiotic stress tolerance syndromes are the main drivers of range filling in temperate woody species. Finally, our results also demonstrate the importance of considering trait syndromes, reflected here as differences between plant functional types, in clarifying the ways in which species’ adaptations influence broad interspecific differences in range size in woody plant species in relation to abiotic stress tolerance.

## Acknowledgements

GP was supported by the Estonian Research Council grant (PSG708), and by the Estonian University of Life Sciences (P200187PKEL). LL was supported by the Estonian University of Life Sciences (P200190PKEL). GP also thanks Babak Naimi for helpful discussion on the implementation of SDMs.

## Notes

### Competing Interest Statement

The authors have declared no competing interest.

