## Appendixes for "The cold-drought tolerance trade-off in temperate woody plants constrains range size, but not range filling": AppendixS1.docx (from another version of this preprint: version 1, 2021-07-25)

**Appendix S1**

**GBIF data cleaning procedure**

The GBIF data were retrieved from multiple sources (**Table S1.1**) using the R function *occ_search* included in the *rgbif* R package (Chamberlain & Boettiger, 2017). The query was set to return a maximum of 20,000 data points per species with associated metadata. Problematic records were identified using the R function *clean_coordinates* in the package *CoordinateCleaner* (Zizka *et al.*, 2020). This function implements several criteria that rely on both species presence data and associated metadata. Automated cleaning was limited to the following criteria: coordinate validity; equal latitude and longitude; zero coordinates; coordinates corresponding to country capitals or centroids; coordinates in the proximity of GBIF headquarter or biodiversity institutions.

The GBIF data were subsequently cleaned by removing duplicates and records derived from fossil specimens, literature, machine observations, or with unknown origin. Since pre-1980 observations are considered unreliable (Maldonado *et al.*, 2015), only records from 1980 onwards were retained. We adopted the latitudinal and longitudinal extent of North America and Europe from the ENVIREM raster files (Title & Bemmels, 2018), so that presence data falling outside the native continent of a species were excluded. To further increase reliability of presence data, we preferred to use a conservative approach by filtering out occurrences outside the polygon defining the actual range of a species (**Fig. S1.1**).

Occurrence data were also subjected to environmental filtering following Varela *et al.* (2014). The *gridSample* function of the R *dismo* package was used for the filtering process (Hijmans *et al.*, 2017). This function selects points from a two-dimensional space using a user-defined grid as stratification. To define the x- and y-axis of the climatic space we implemented a Principal Component Analysis (PCA) using the environmental layers presented in the main text. The first and second PCA axes were used as the x‐ and y‐axis defining the environmental space. Repetitive clumped points in this space were filtered out to maximize the evenness of occurrence data climatic coverage (Varela *et al.*, 2014) (**Fig. S1.1**). This procedure was carried out separately for each species. Each generated climatic grid had the same resolution as the environmental layers so that only one occurrence per grid cell was retained (**Fig. S1.1**). Environmentally filtered data were further checked for outliers and values above the 95^th^ percentile were omitted (Nogués-Bravo *et al.*, 2014; Estrada *et al.*, 2015).

**Fig. S1.1.** Representation of the filtering process applied to obtain reliable presence data to fit species distribution models. 1: filtering of species presence data obtained from the Global Biodiversity Information Facility (GBIF) to keep only data points falling within each species’ actual range; and 2: their subsequent environmental filter (light blue area). All the example phases include information for *Abies alba* only, but the same procedure was applied to each of the 331 woody plant species included in the study. The grids representing the environmental space and the reduction of data points clumping have only demonstrative purposes and are not based on real data. The polygon describing *A. alba*’s actual range was obtained from European forest genetic resources program (EUFORGEN, <http://www.euforgen.org/species>).

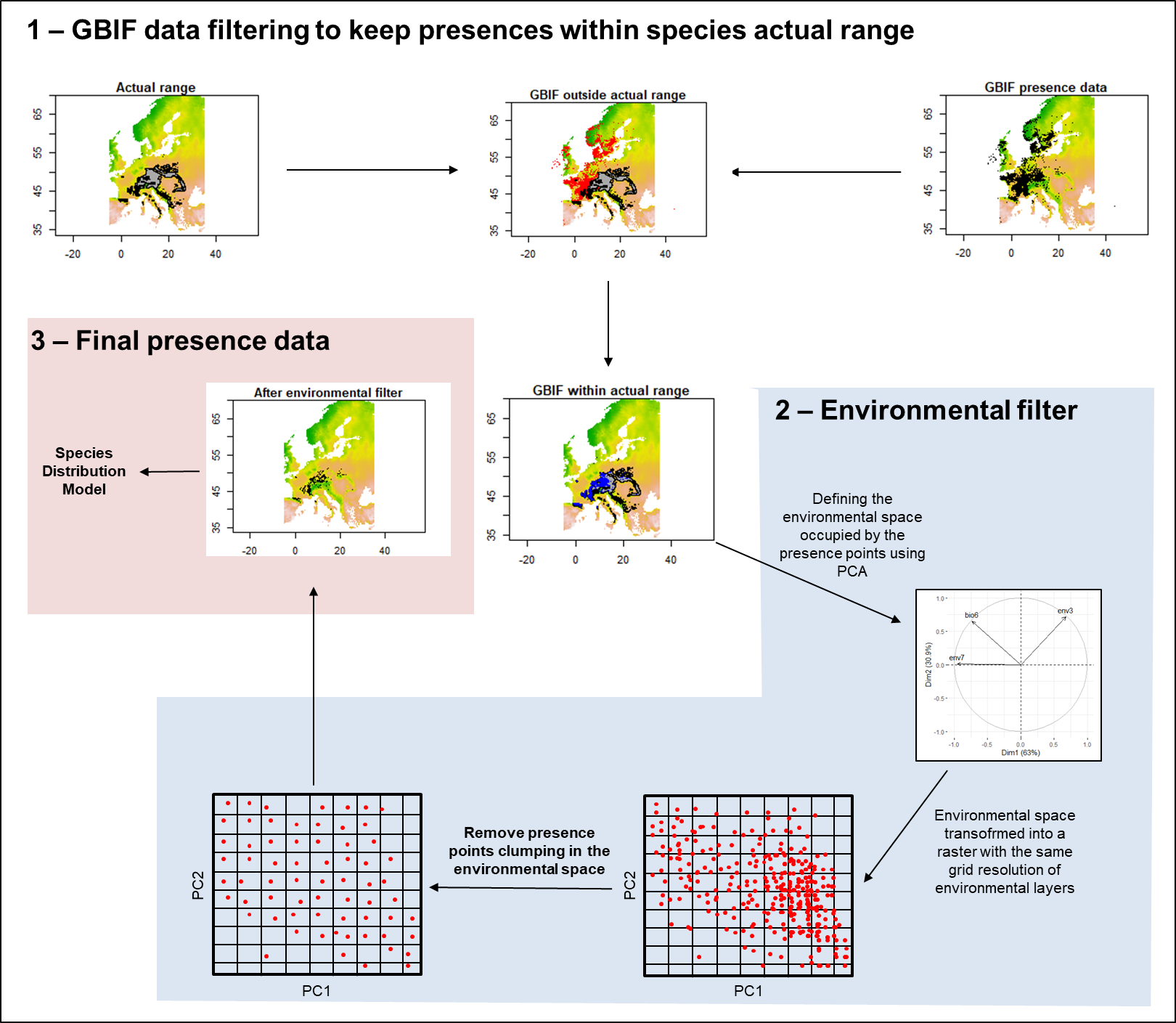
