## Appendixes for "The cold-drought tolerance trade-off in temperate woody plants constrains range size, but not range filling": AppendixS2.docx (from another version of this preprint: version 1, 2021-07-25)

**Appendix S2**

**Example of range filling calculation**

**Fig. S2.2.** Example of range filling calculation. The extent of species actual range (number in the top right portion of the Species actual range panel) was calculated as the number of raster cells occupied by the polygon drawn around each species’ actual range (black area). The obtained actual range value (i.e. 2201) was divided per each potential range size estimate (number in the top right portion of the panels) obtained using the Bioclim algorithm (green areas, see Material and Methods) and calculated as the number of green cells in the maps. The black line in each potential range size panel shows the overlap between species actual and potential range. For demonstrative purposes only, calculations were reported here only reported here for four replicates but a total of 30 replicates per species were used to estimate a single mean range filling value. All the information included in the figure refers to *Abies alba*. The polygon describing *A. alba* actual range was obtained from European forest genetic resources program (EUFORGEN, <http://www.euforgen.org/species>).

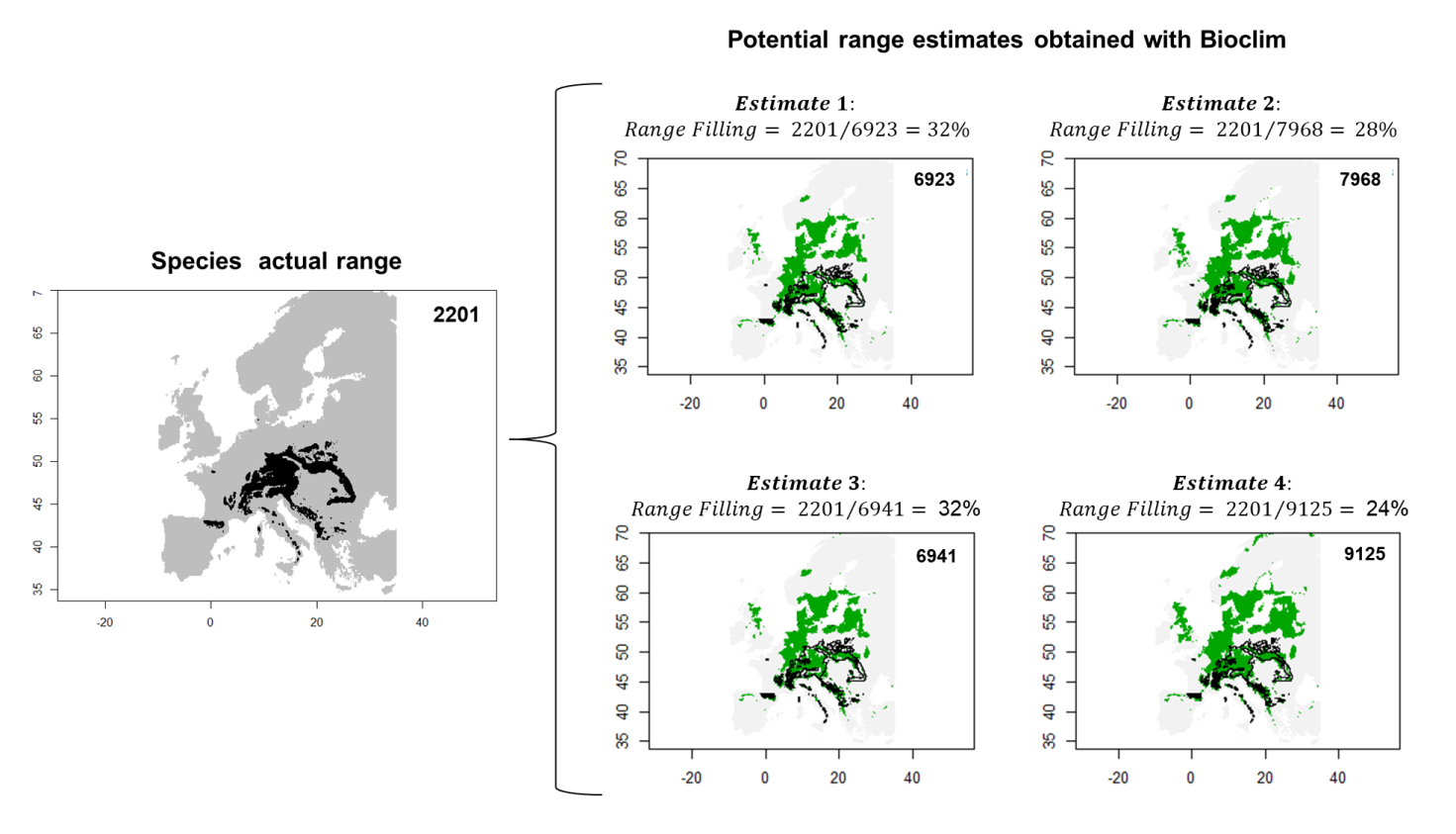
