## Appendixes for "The cold-drought tolerance trade-off in temperate woody plants constrains range size, but not range filling": AppendixS3.docx (from another version of this preprint: version 1, 2021-07-25)

**Appendix S3**

**Species Distribution Models output, performance and considerations on range filling estimates**

Estimates of either potential range or range filling obtained from Bioclim and Maxent were highly correlated (**Fig. S3.3**). However, range filling calculated using Bioclim derived estimates of potential range had fewer values > 100% (18% vs. 50% in Bioclim vs. Maxent, respectively), therefore only results based on the Bioclim derived estimates were considered for further analyses. Bioclim performance was overall good, with most TSS and AUC values above 0.80 (**Fig. S3.4**). The correlation coefficient frequency for GDD, CMI and T_min_ is shown in **Fig. S3.5**. Our actual range and range filling estimates were highly correlated to those reported by the literature for a similar species set (**Fig. S3.6 a,b**).

The species displaying range filling estimates > 100% based on the Bioclim datawere either very widely or very narrowly distributed species (**Table S3.1**), and they may therefore reflect the limitations of the model for predictions where species distributions are somewhat extreme. For example, in some cases, the native distribution of some species extended further than the studied continent boundaries, as in the case of Eurasian species (**Table S3.1**). At any rate, a similar proportion (17%) of species with range filling values > 100% was also obtained in a previous study including widely distributed European tree species (Nogués-Bravo *et al.*, 2014).

Some models showed poor accuracy (i.e. TSS < 0.6) but, again, they were for species with range filling values > 100% (**Fig. S3.2**). Also, correlation coefficients with the selected environmental predictors tend to be lower for the models relative to species with a range filling > 100% (**Fig. S3.3**), possibly reflecting that their wide distribution is not constrained by the considered environmental variables.

Specific methodological choices could have also affected our range filling estimates. We filtered out occurrence points falling outside the known range of the species but the retained occurrences may not cover the entire geographical area and/or range of climatic conditions experienced by a species. As a result, potential range sizes can be underestimated and range filling overestimated, but this limitation applies to all species, not only the ones with range filling above 100%. However, since our data included European and North Americana species, and presence data for species from these continents are known to be more reliable compared to other parts of the world (Zizka *et al.*, 2020), we still consider the potential range size estimates based on presence data only reliable.

At any rate, we did not exclude values above 100% from the analysis because they are representative of high range filling for very widely or very narrowly distributed species. Our range filling estimates using the same algorithm, very similar environmental layers and a similar species set, are in fact highly correlated with those from Svenning & Skov (2004), Nogués-Bravo *et al.* (2014) and Seliger *et al.* (2021) (**Fig. S3.6**) and qualitatively comparable despite the different resolution of environmental layers. A very high correlation was also found between our actual range size estimates (log_10_-transofrmed) and those reported by Svenning & Skov (2004), Nogués-Bravo *et al.* (2014) and Seliger *et al.* (2021) (**Fig. S3.6**).

**Fig. S3.3.** Correlation between species-specific **(a):** mean potential range size (n cells, see Materials and Methods) estimated using Bioclim and Maxent algorithms; and **(b):** mean range filling estimates (n cells) derived from Bioclim and Maxent. Correlation coefficients (R) are shown (n = 331). Each data point is the mean of 30 replicates.

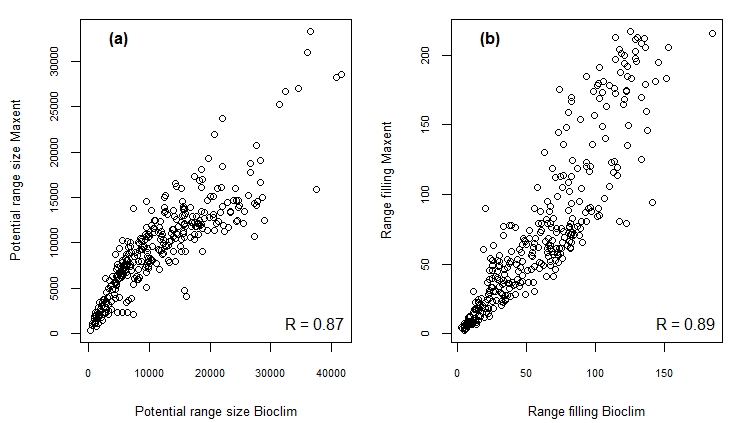

**Fig. S3.4.** Frequency distribution of mean values of True Skill Statistic (TSS) and Area Under the Receiving Operator Curve (AUC) for 331 species distribution models (each value was the mean of 30 replicates per species). The first row shows TSS and AUC frequency distributions for all the models. The second and the third rows display TSS and AUC frequency distributions for the models from which we derived species-specific range filling estimates above and below 100%.

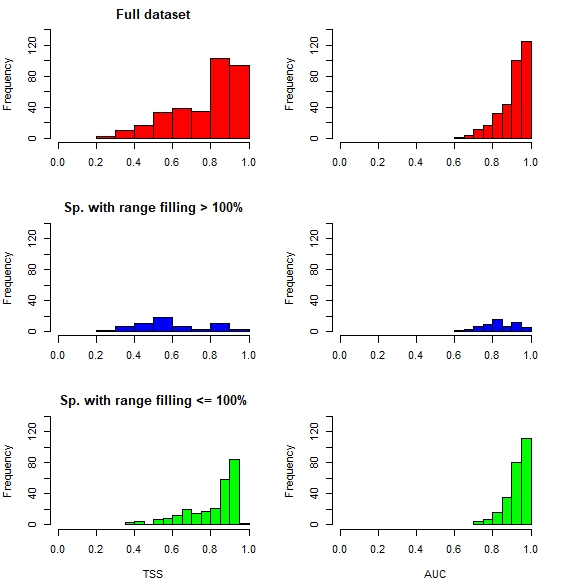

**Fig. S3.5.** Frequency distributions of mean values of correlation coefficients for the considered environmental layers (Tmin = mean minimum temperature of the coldest month; GDD = Growing degree days above 5°C; CMI = climatic moisture index; see Materials and Methods for a detailed definition of the environmental layers) for 331 species distribution models (each value was the mean of 30 replicates per species). The first row shows the frequency distribution of correlation coefficient values for all the models run in the present study. The second and the third rows show frequency distributions of correlation coefficient values for the models from which we derived species-specific range filling estimates above and below 100%.

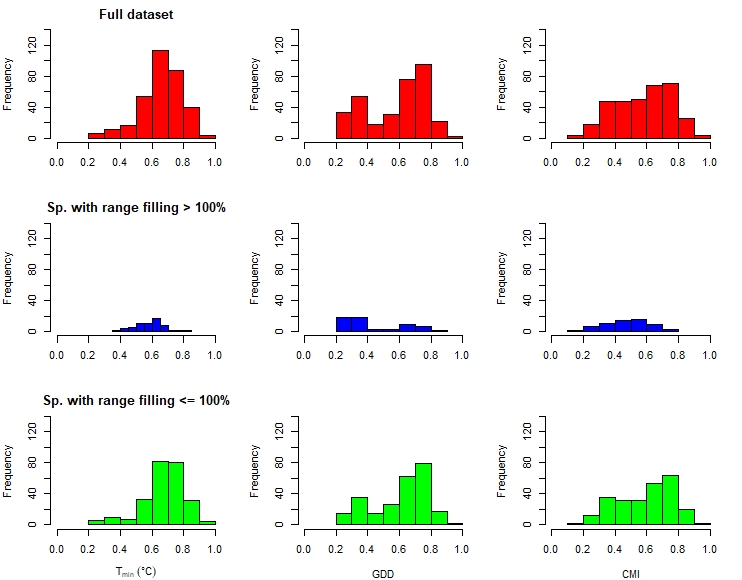

**Fig. S3.6.** Relationships between: **(a)** actual range size and **(b)** range filling estimates obtained in our study with those reported by the literature. Data from the literature (n = 225) were gathered from Svenning & Skov (2004), Nogués-Bravo *et al.* (2014) and Seliger et al. (2021) and span both European and North American species. Slope, correlation coefficient (R) and p value for the Ordinary Least Square (OLS) fits are shown. Solid black lines depict quantile regressions, which were always significant at p ˂ 0.05. The blue and the red lines in panel **(b)** refer to the OLS fit obtained by including or excluding range filling estimates above 100%, respectively (see **Supporting Note 2**).

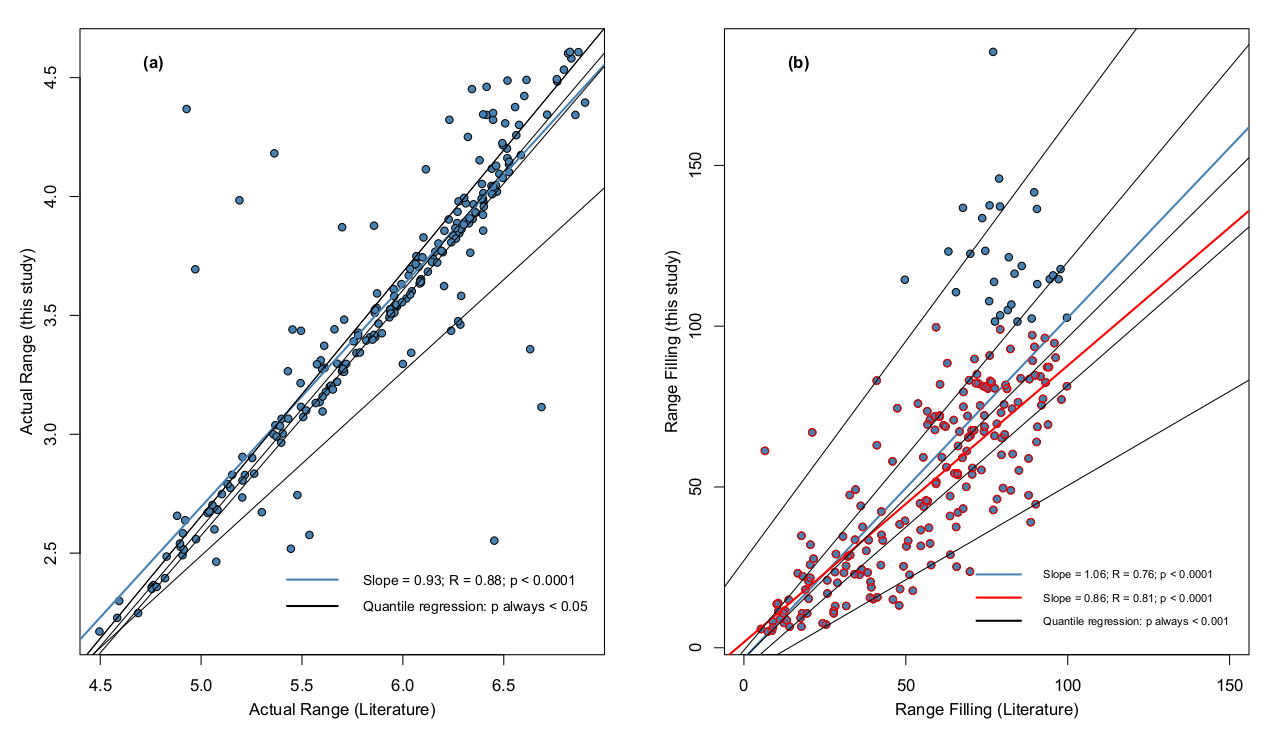
