## Appendixes for "The cold-drought tolerance trade-off in temperate woody plants constrains range size, but not range filling": AppendixS5.docx (from another version of this preprint: version 1, 2021-07-25)

**Appendix S5**

**Table S5.3.** Phylogenetic Generalized Linear Mixed Models run on all data pooled. Models were run using Actual range (log_10_-transformed number of 10 arcmin cells) and range filling (Residuals, see Methods) as response variables, Stress Axis 1 (Standard Deviation units) and Latitude (centroid latitude, °) as main effects, and Plant Functional Type (*PFT*) and species phylogenetic relatedness (*Phylogeny*) as random effects. Estimates of the main effects and their standard errors (SE) together with the model explained variance (R^2^), and sample size are shown. The variance of the response variable captured by the random effects is also shown. Significance code: *** indicates p ≤ 0.0001; ** p ≤ 0.001; * p ≤ 0.01; ns = not significant.

| **Response** | **Main effect** | |  |  |  |  | **Random effect** | |
| --- | --- | --- | --- | --- | --- | --- | --- | --- |
|  | **Predictors** | **Slope** | **SE** | **p** | **R^2^** |  |  | **Variance** |
| Actual range | *Intercept* | 2.770 | 0.180 | *** | 0.22 |  | *PFT* | 0.03 |
|  | *Stress Axis* | -0.08 | 0.030 | ** |  |  | *Phylogeny* | ≈ 0 |
|  | *Latitude* | 0.010 | 0.003 | *** |  |  |  |  |
| Range filling (Residuals) | *Intercept* | 20.060 | 7.810 | * | 0.04 |  | *PFT* | 8.5 |
|  | *Stress Axis* | -3.350 | 1.270 | ** |  |  | *Phylogeny* | 0.001 |
|  | *Latitude* | -0.490 | 0.170 | ** |  |  |  |  |

**Table S5.4.** Phylogenetic Generalized Linear Mixed Models run using only species belonging to the genera shared between continents. Models were run using Actual range (log_10_-transformed number of 10 arcmin cells) and range filling (Residuals, see Methods) as response variables, Stress Axis 1 (Standard Deviation units) and Latitude (centroid latitude, °) as main effects, and Plant Functional Type (*PFT*) and species phylogenetic relatedness (*Phylogeny*) as random effects. Estimates of the main effects and their standard errors (SE) together with the model explained variance (R^2^), and sample size are shown. The variance of the response variable captured by the random effects is also shown. Significance code: *** indicates p ≤ 0.0001; ** p ≤ 0.001; * p ≤ 0.01; ns = not significant.

|  |  | **Main effects** |  |  |  |  |  |  | **Random effects** |  |
| --- | --- | --- | --- | --- | --- | --- | --- | --- | --- | --- |
| **Response** | **Continent** |  | **Estimate** | **SE** | ***p*** | ***R*^2^** | **n** |  |  | **Variance** |
| **Actual range** | **Europe** | *Intercept* | 3.670 | 0.408 | *** | 0.12 | 92 |  | *PFT* | ≈ 0 |
|  |  | *Stress Axis 1* | -0.150 | 0.049 | ** |  |  |  | *Phylogeny* | ≈ 0 |
|  |  | *Latitude* | 0.0001 | 0.008 | ns |  |  |  |  |  |
|  | **North America** | *Intercept* | 2.847 | 0.263 | *** | 0.27 | 148 |  | *PFT* | ≈ 0 |
|  |  | *Stress Axis 1* | -0.120 | 0.038 | ** |  |  |  | *Phylogeny* | ≈ 0 |
|  |  | *Latitude* | 0.012 | 0.006 | * |  |  |  |  |  |
| **Range filling** | **Europe** | *Intercept* | -1.899 | 19.206 | ns | 0.21 | 92 |  | *PFT* | ≈ 0 |
|  |  | *Stress Axis 1* | -53.250 | 12.922 | *** |  |  |  | *Phylogeny* | ≈ 0 |
|  |  | *Latitude* | 0.123 | 0.398 | ns |  |  |  |  |  |
|  |  | *Stress Axis 1 * Latitude* | 0.900 | 0.249 | ** |  |  |  |  |  |
|  | **North America** | *Intercept* | 32.636 | 11.689 | * | 0.06 | 148 |  | *PFT* | ≈ 0 |
|  |  | *Stress Axis 1* | -0.477 | 1.875 | ns |  |  |  | *Phylogeny* | ≈ 0 |
|  |  | *Latitude* | -0.822 | 0.284 | * |  |  |  |  |  |

**Fig. S5.7.** Relationship between the cold-drought trade-off (Stress Axis 1, Standard Deviation units), Centroid latitude (°) and actual range (log10-transformed number of 10 arcmin cells) and range filling (residuals, see Methods) for: (a,b) European (n = 128), and (c, d) North American (n = 197) woody species. The color gradient from blue to red reflects the progression from cold/wet tolerant to warm/dry tolerant species along the cold-drought trade-off (i.e. Stress Axis 1). Statistics are reported in **Table 1** in the main text.

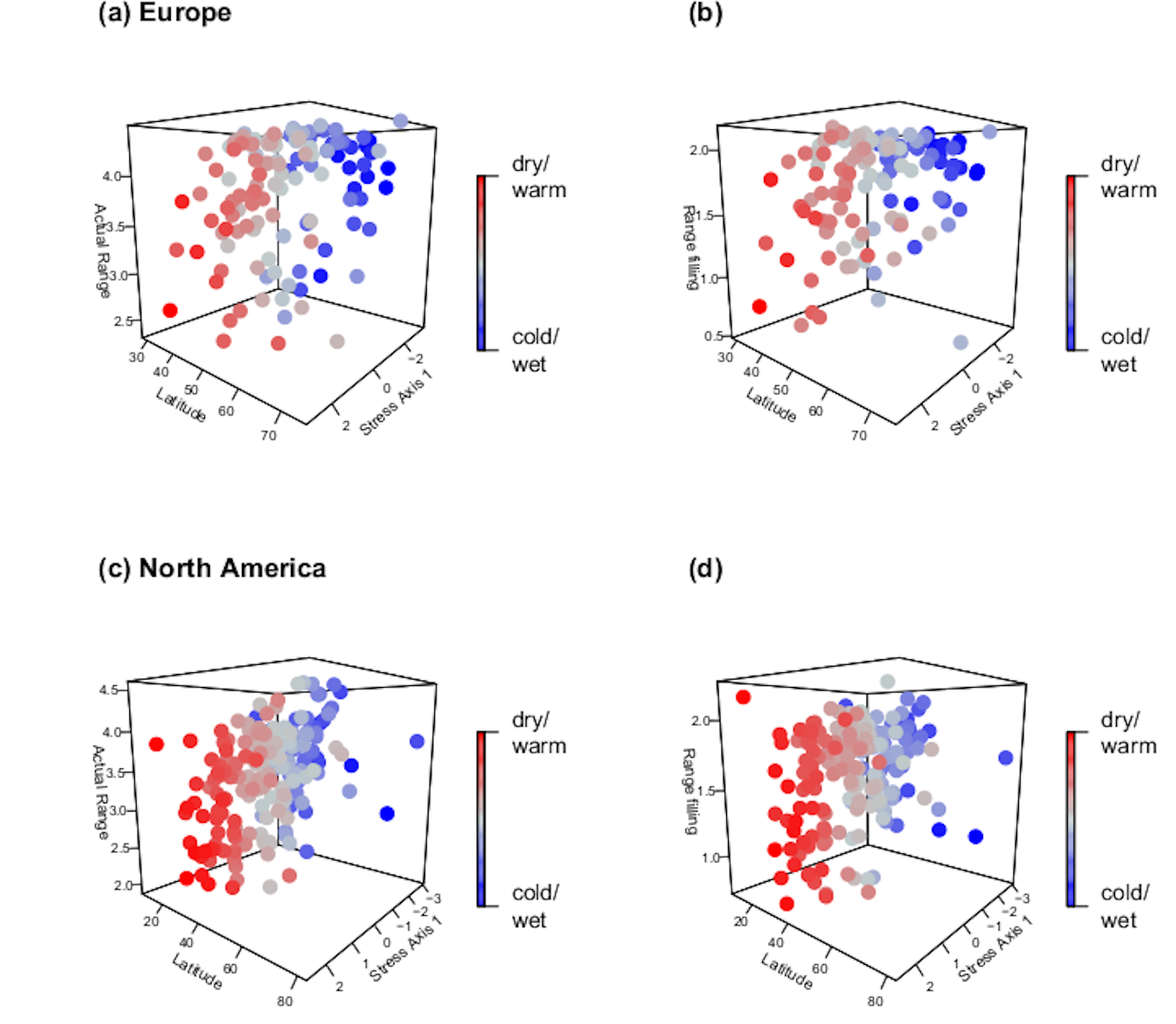
