## Appendixes for "The cold-drought tolerance trade-off in temperate woody plants constrains range size, but not range filling": AppendixS6.docx (from another version of this preprint: version 1, 2021-07-25)

**Appendix S6**

**Results of the spatial autocorrelation test for the residuals of the Phylogenetic Generalized Linear Mixed Models (PGLMMs).**

**Fig. S6.8.** Spline correlograms showing the degree of spatial autocorrelation in Phylogenetic Generalized Linear Mixed Models residuals. Panel headings show the response variable – continent combination. The full model structure is shown in Table 1 in the main text. Shaded areas around the mean estimates (solid black line) represent 95% pointwise bootstrap confidence intervals computed by 5,000 bootstrap samples of Pearson’s residuals. (a) Actual range for European continent, (b), actual range for North American continent, (c) Range filling in Europe, and (d) range filling in North America.

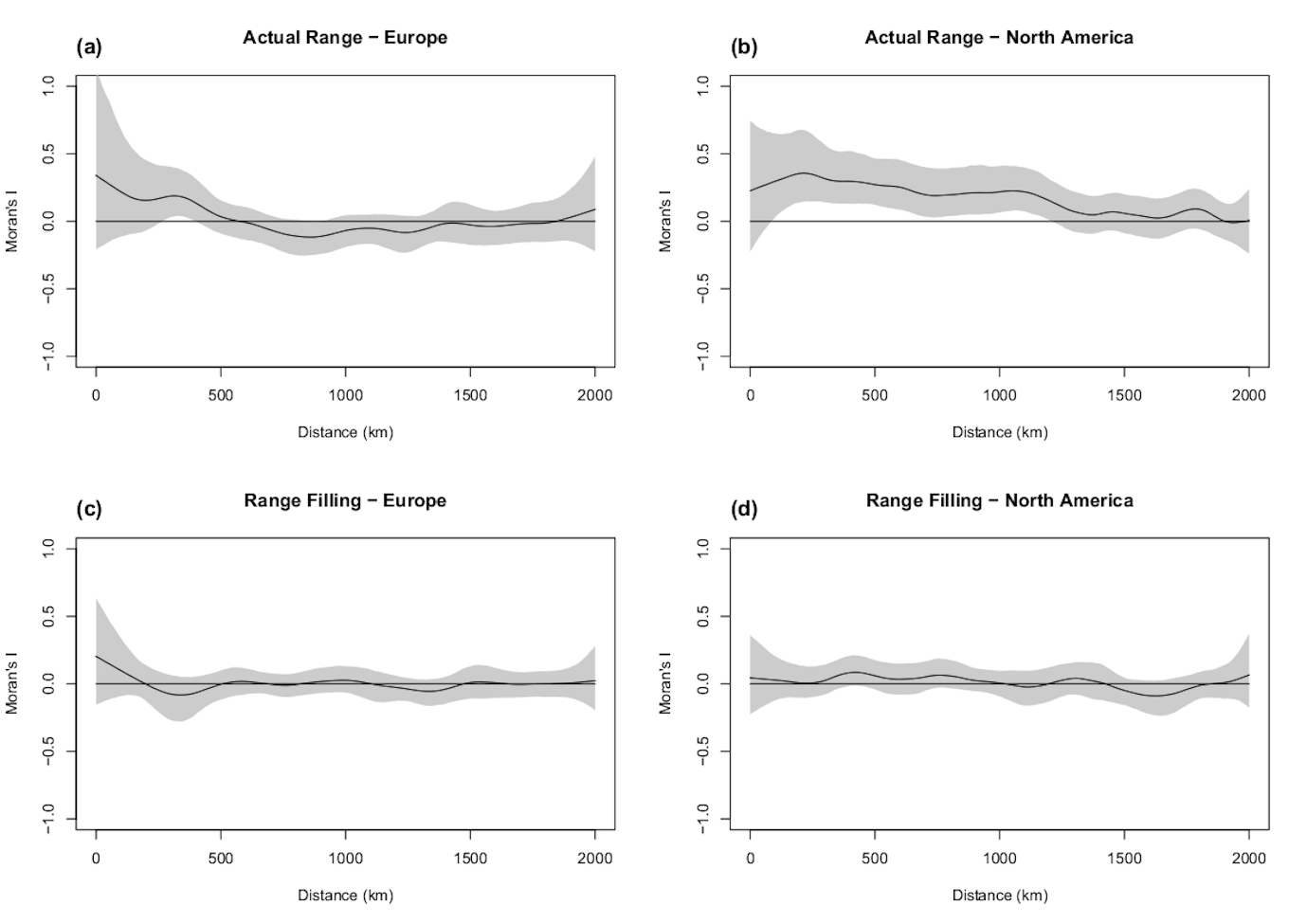
